# Structural analysis of the macrocyclic inhibitor BI-4020 binding to EGFR kinase

**DOI:** 10.1101/2022.08.27.505540

**Authors:** Tyler S. Beyett, Jaimin K. Rana, Ilse K. Schaeffner, David E. Heppner, Michael J. Eck

## Abstract

A novel macrocyclic inhibitor of mutant EGFR (**BI-4020**) has shown promise in pre-clinical studies of T790M and C797S drug-resistant non-small cell lung cancer. To better understand the molecular basis for BI-4020 selectivity and potency, we have carried out biochemical activity assays and structural analysis with X-ray crystallography. Biochemical potencies agree with previous studies indicating that BI-4020 is uniquely potent against drug-resistant L858R/T790M and L858R/T790M/C797S variants. Structures show that BI-4020 is likely rendered selective due to interactions with the kinase domain hinge region as well as T790M, akin to Osimertinib. Additionally, BI-4020 is also rendered more potent due to its constrained macrocycle geometry as well as additional H-bonds to conserved K745 and T845 residues in both active and inactive conformations. These findings taken together show how this novel macrocyclic inhibitor is both highly potent and selective for mutant EGFR in a reversible mechanism and motivate structure-inspired approaches to developing targeted therapies in medicinal oncology.

---

Mutations in the epidermal growth factor receptor (EGFR) are common oncogenic drivers of non-small cell lung cancer (NSCLC). The most common oncogenic point mutation in the EGFR kinase domain is L858R, which increases kinase enzymatic activity and K_m,ATP_, thereby rendering the kinase highly susceptible to ATP-competitive tyrosine kinase inhibitors (TKIs).^1,2^ Patients treated with the FDA-approved 1^st^-generation TKIs gefitinib (Figure 1) or erlotinib almost invariably acquire the T790M resistance mutation at the gatekeeper position, which restores the K_m,ATP_ of the L858R variant to that of the wild-type (WT) kinase thereby closing the therapeutic window.^3^ To overcome acquired resistance from T790M, subsequent inhibitor generations incorporated an acrylamide warhead that covalently bonds with C797.^4,5^

While the lack of mutant selectivity in 2^nd^-generation inhibitors such as afatinib limited their use, the development and clinical approval of the mutant-selective 3^rd^-generation irreversible inhibitor osimertinib has greatly changed the treatment landscape.^6,7^ However, patients with NSCLC treated with osimertinib ultimately develop drug resistance, including through acquisition of an additional third mutation, C797S, which abolishes drug efficacy by preventing the formation of the potency-enabling covalent bond.^8–10^

**Figure 1:**
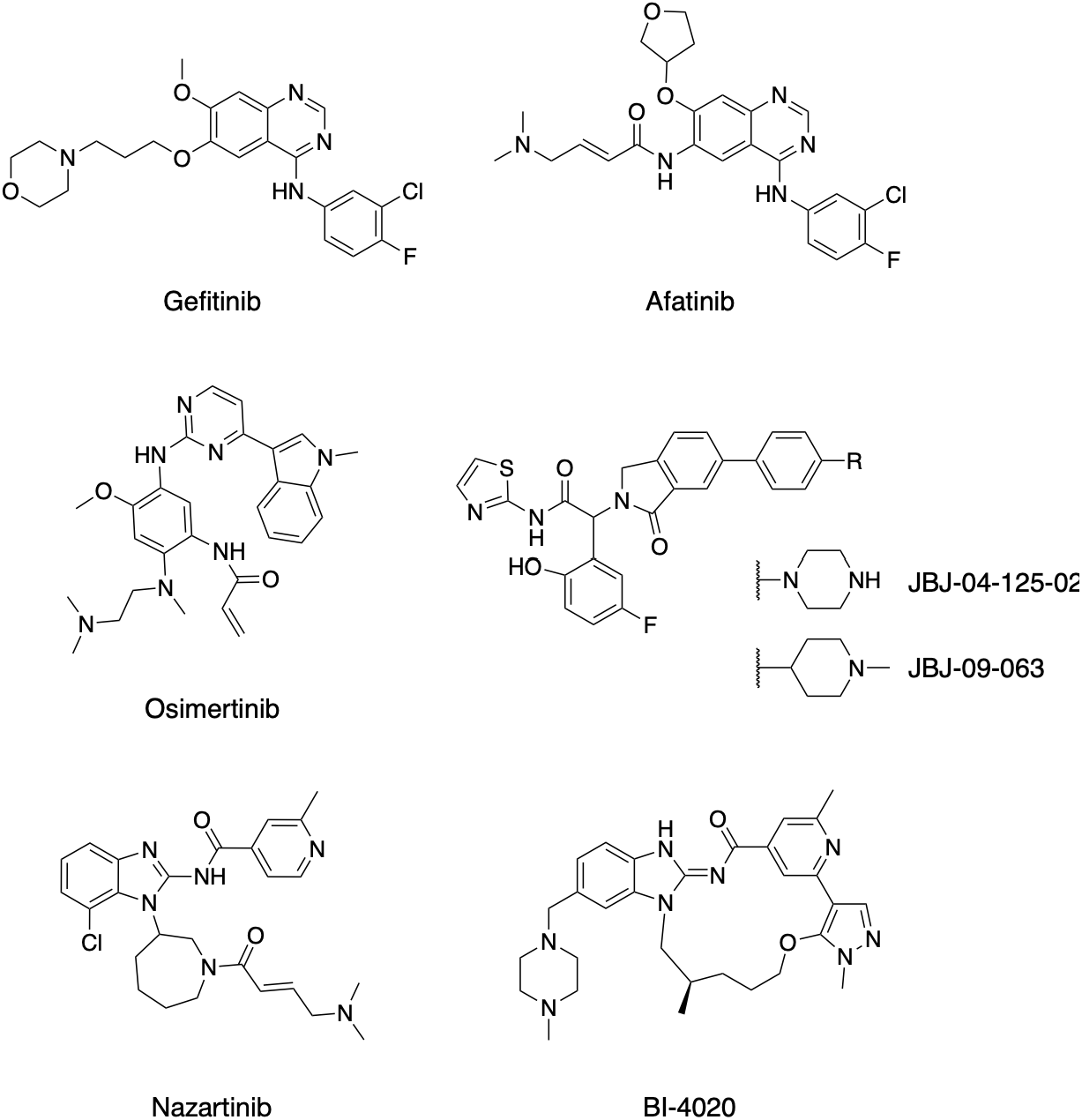
Chemical structures of EGFR inhibitors.

Extensive efforts to develop inhibitors against L858R/T790M/C797S drug-resistant NSCLC have generated various compounds targeting the ATP substrate site^11–16^ as well as mutant-selective allosteric inhibitors, such as JBJ-04-125-02 and JBJ-09-063 (Figure 1).^17–20^ A key attribute of the recently emerged next-generation L858R/T790M/C797S ATP-competitive tool compounds is their ability to afford stronger reversible binding to the kinase domain through interactions that often come at the cost of drug selectivity for the oncogenic mutant.^21^ The macrocyclic ATP-competitive inhibitor **BI-4020** is structurally related to the 3^rd^-generation inhibitor nazartinib but displays exquisite potency against drug-resistant L858R/T790M/C797S and del19/T790M/C797S variants through a reversible binding mode.^22^ The initial report of **BI-4020** described its efficacy in a variety of cell lines and tumor models *in vivo* and reported crystal structures of precursor molecules.^22^ We build upon this work by reporting enzymatic potencies of **BI-4020** against EGFR variants and HER2 and through the determination of crystal structures of **BI-4020** in complex with EGFR in the active and inactive conformations.

To better understand the molecular nature of **BI-4020** efficacy, we conducted enzymatic inhibition assays using recombinant EGFR kinase domain variants and the related HER2 kinase domain. For these purposes, we utilized the HTRF KinEASE assay system along with purified EGFR kinase domain, including the L858R/C797S mutant, which is highly relevant for front-line osimertinib resistance.^8,9^ To obtain the most comprehensive set of biochemical potencies, we compared **BI-4020** to the closely related non-macrocyclic inhibitor nazartinib as well as to gefitinib, osimertinib, and mutant-selective allosteric inhibitor JBJ-04-125-02.^17^ **BI-4020** was found to be less potent than the others at inhibiting EGFR(L858R) but was more potent (IC_50_=0.01 nM) and selective for EGFR(L858R/T790M) and EGFR(L858R/T790M/C797S). As expected, the 1^st^-generation reversible TKI gefitinib is ineffective against T790M variants, and both irreversible inhibitors (osimertinib and nazartinib) were poor inhibitors of C797S-containing variants. The allosteric inhibitor JBJ-04-125-02 was the only other inhibitor assayed that was effective at inhibiting C797S-containing variants. Similar to nazartinib, **BI-4020** was not a potent inhibitor of the related HER2 kinase. Consistent with prior work ^22^, these biochemical studies demonstrate that **BI-4020** is an exquisitely potent inhibitor of L858R/T790M and L858R/T790M/C797S variants of EGFR, with relative sparing of wild-type (WT) EGFR and HER2.

**Table 1:**
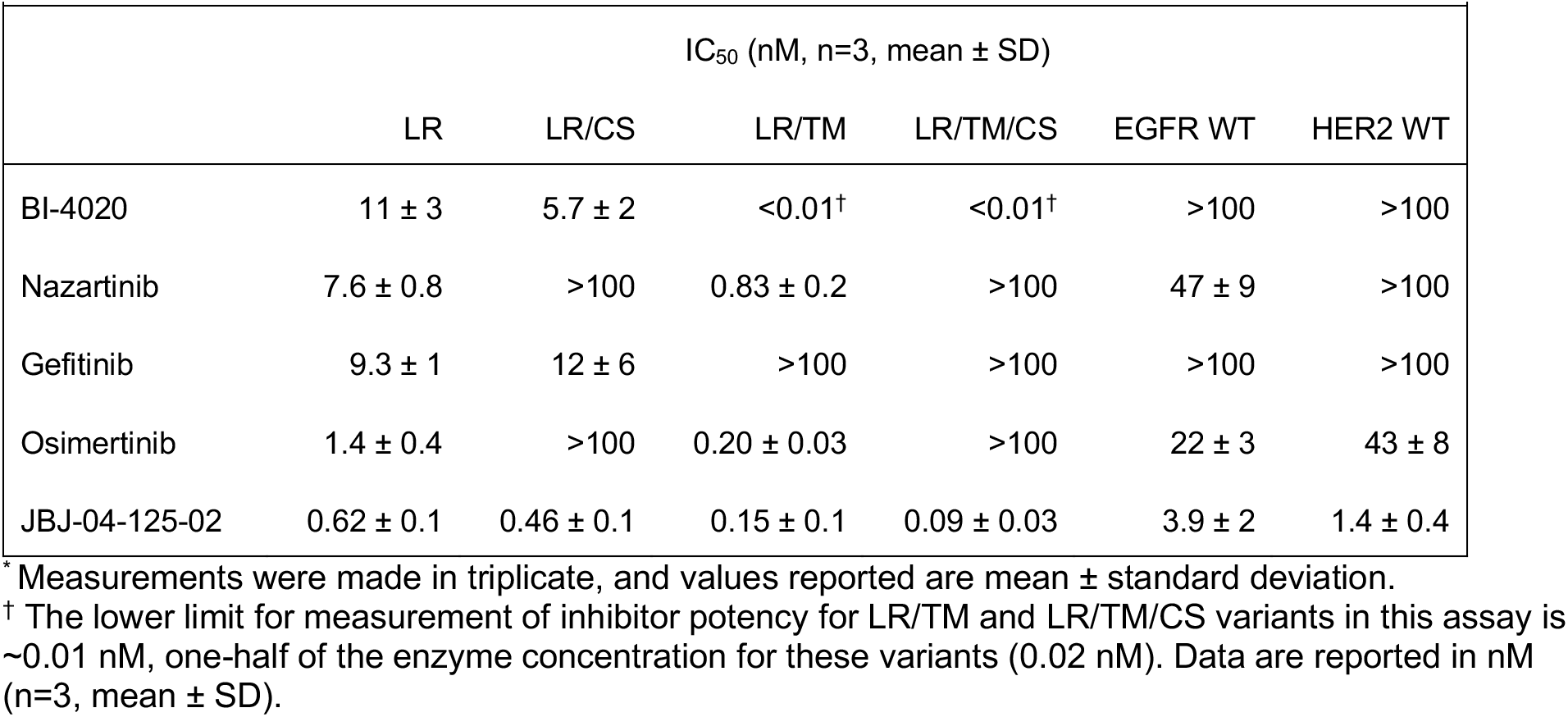
Biochemical potency of BI-4020 against selected EGFR variants and HER2.*

|  |  |  |  |  |  |  |
| --- | --- | --- | --- | --- | --- | --- |
|  | IC<sub>50</sub> (nM, n=3, mean ± SD) |  |  |  |  |  |
|  | LR | LR/CS | LR/TM | LR/TM/CS | EGFR WT | HER2 WT |
| BI-4020 | 11 ± 3 | 5.7 ± 2 | <0.01† | <0.01† | >100 | >100 |
| Nazartinib | 7.6 ± 0.8 | >100 | 0.83 ± 0.2 | >100 | 47 ± 9 | >100 |
| Gefitinib | 9.3 ± 1 | 12 ± 6 | >100 | >100 | >100 | >100 |
| Osimertinib | 1.4 ± 0.4 | >100 | 0.20 ± 0.03 | >100 | 22 ± 3 | 43 ± 8 |
| JBJ-04-125-02 | 0.62 ± 0.1 | 0.46 ± 0.1 | 0.15 ± 0.1 | 0.09 ± 0.03 | 3.9 ± 2 | 1.4 ± 0.4 |

\* Measurements were made in triplicate, and values reported are mean ± standard deviation.
† The lower limit for measurement of inhibitor potency for LR/TM and LR/TM/CS variants in this assay is ~0.01 nM, one-half of the enzyme concentration for these variants (0.02 nM). Data are reported in nM (n=3, mean ± SD).

To reveal the binding mode of **BI-4020** and better understand its selectivity, we determined co-crystal structures in complex with EGFR kinase domain in both active and inactive conformations at 2.4 Å and 3.1 Å resolution, respectively (Figure 2, SI Table 1, SI Figure 1). We employed WT EGFR kinase for the active-state structure with **BI-4020,** and the T790M/V948R variant for the inactive state (the V948R mutation in the kinase C-lobe disrupts formation of the activating dimer interaction, allowing crystallization in the inactive, αC-helix out, conformation). In both structures, **BI-4020** interacts with the kinase hinge through the amide carbonyl and benzimidazolidine, similarly to the binding mode of nazartinib (Figure 2C).^23^ In our structure with T790M, favorable hydrophobic contacts between the side chain and aromatic ring of BI-4020 and a stacking interaction between the aromatic pyridine and methionine (Figure 2D). Similar contacts have been shown through a crystal structure and molecular dynamics to be important for the T790M selectivity of osimertinib indicating that this is the likely mechanism where **BI-4020** acquires selectivity of the T790M mutations (SI Figure 2A, B).^24^ These structures provide additional examples of macrocyclic EGFR inhibitors, of which there are few reported.^22,25^ These structures are useful for structure-based drug design, as rigidification is a well-established strategy for reducing the entropic cost associated with ligand binding and can also improve other pharmacokinetic (PK) properties such as metabolic stability, cellular permeability, and oral bioavailability.^26^

**Figure 2:**
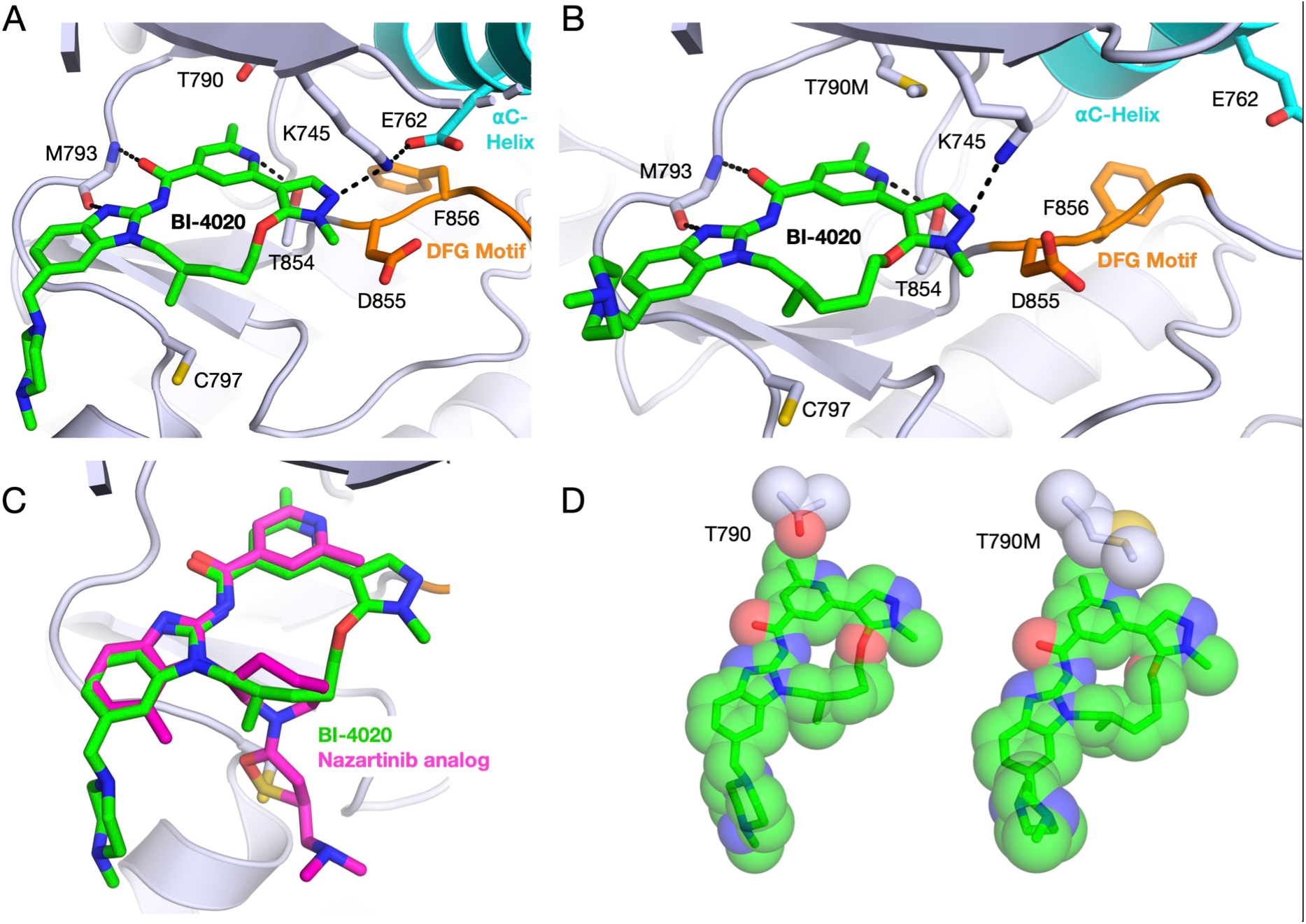
Crystal structures of EGFR bound to **BI-4020**. A) **BI-4020** in complex with WT EGFR in the active conformation (PDB 7KXZ). In this conformation, E762 is positioned towards the ATP site and interacts with the catalytic K745. B) **BI-4020** in complex with EGFR(T790M/V948R) in the inactive conformation (PDB 7KY0). E762 is positioned away from the active site, and K745 still interacts with the inhibitor. C) Overlay of WT EGFR in complex with **BI-4020** (green) or a close analog of nazartinib (magenta, PDB 5FEQ). D) Comparison of **BI-4020** contacts with the gatekeeper residue 790. Limited hydrophobic contacts and a possibly disfavored contact between the T790 side chain hydroxyl and **BI-4020** methyl are observed. In contrast, more extensive hydrophobic contacts and a favorable sulfur-aromatic stacking interaction are observed in the T790M variant.

The necessity of forming covalent bonds to C797 by highly-effective 3^rd^-generation TKIs has spurred significant efforts in discovering molecules that can inhibit C797S-containing variants as the next generation of EGFR inhibitors. An earlier structural study from our lab characterized the mechanism by which trisubstituted imidazole inhibitors acquire enhanced reversible binding through potency-enabling H-bonding to K745 in the inactive conformation (Figure 3A, B, SI Figure 3).^21^ Analogous H-bonds to K745 are made by **BI-4020,** regardless of C-helix conformation, and several other C797S-active inhibitors including CH7233163, JN3229, brigatinib derivatives, and others.^13,14,21,25,27^ An additional H-bond with T854 is also common, and the necessity for these H-bonds appears to be a general feature of potent fourth-generation ATP-competitive inhibitors of EGFR(L858R/T790M/C797S).^14,22,27^ While allosteric inhibitors such as JBJ-04-125-02 and JBJ-09-063 are also effective against C797S variants, they cannot cobind with BI-4020 as they can with other TKIs like osimertinib (SI Figure 2C).^28^ However, combination of gefitinib and allosteric inhibitors is beneficial despite a lack of co-binding, suggesting that combinations with **BI-4020** may be of value.^29^

**Figure 3:**
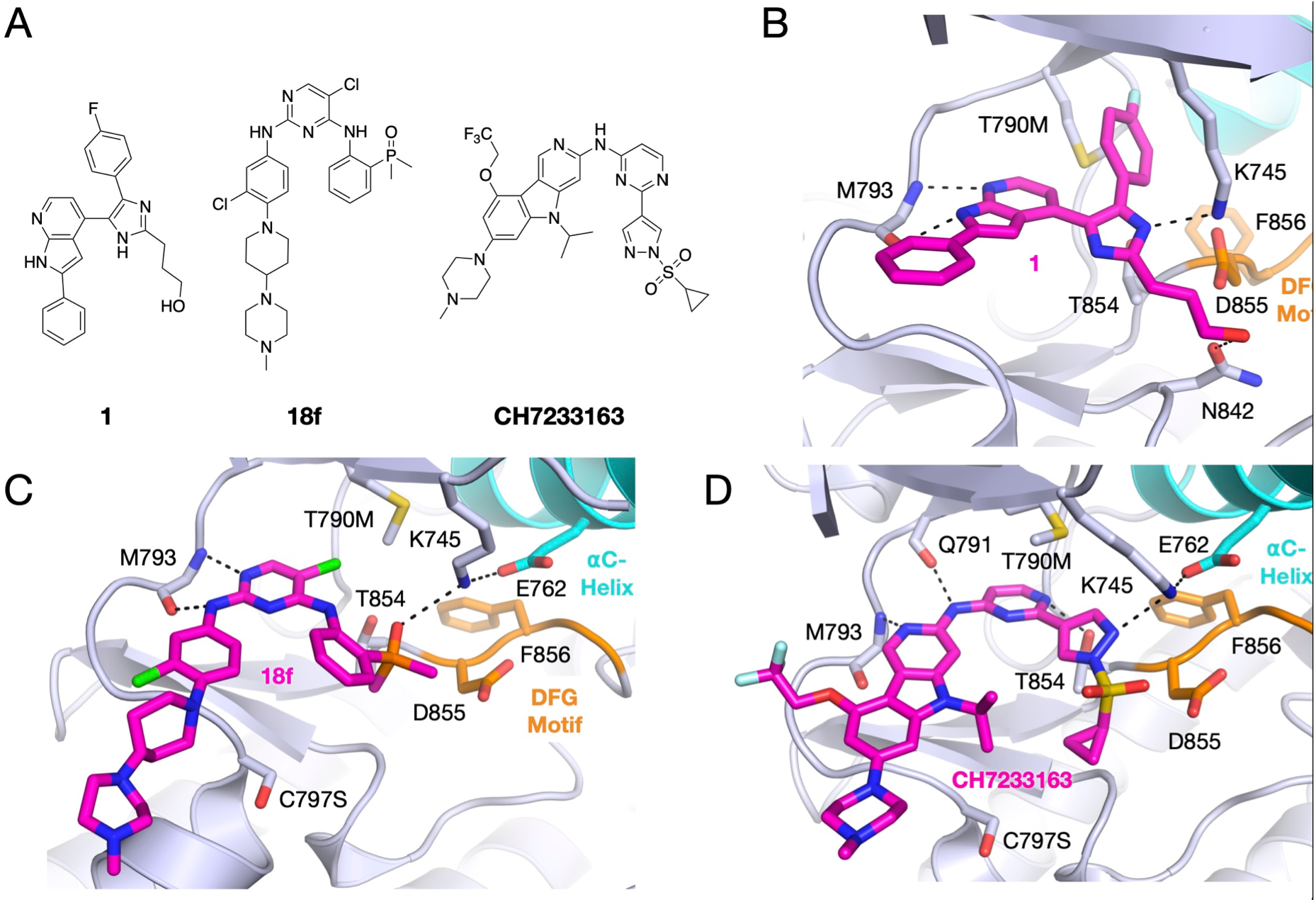
Crystal structures with inhibitors highlighting key strategies for improving reversible binding and mutant selectivity. A) Chemical structures of key inhibitors. B) Trisubstituted imidazole compound LN2084 (**1**) in complex with EGFR(T790M/V948R) in the inactive conformation (PDB 6V5N). C) Brigatinib analog **18f** bound to EGFR(T790M/C797S) in the active conformation (PDB 7ER2). D) CH7233163 in complex with EGFR(L858R/T790M/C797S) (PDB 6LUB).

In conclusion, we have conducted a structural and functional analysis of the macrocycle **BI-4020** as a mutant-selective EGFR inhibitor. Our molecular analysis supports previous biological and medicinal chemistry studies that **BI-4020** is a potent inhibitor of EGFR activity in NSCLC harboring exquisite selectivity for T790M-containing variants, including drug-resistant C797S. **BI-4020** is made potent through multiple strong intermolecular H-bonds afforded by the rigid binding mode enabled due to the macrocyclic nature of the compound. The distinctive selectivity for T790M is made by productive interactions with the gatekeeper methionine, which are less compatible with T790-containing variants. Thus, macrocyclization of existing EGFR inhibitors and engagement of the catalytic K745 appear viable strategies for improving reversible binding and ability to inhibit C797S in next-generation EGFR inhibitors.

## Acknowledgements

We thank the lab of Nathanael Gray for supplying BI-4020 for initial experiments. This work was supported by National Institutes of Health grant RO1 CA201049 (M.J.E.), PO1 CA154303 (M.J.E.), and R35 CA242461 (M.J.E.). T.S.B is supported by a Ruth L. Kirschstein National Research Service Award (F32CA247198-01). D.E.H. acknowledges startup funds from the University at Buffalo, The State University of New York. Research reported in this publication was supported by the National Center for Advancing Translational Sciences of the NIH under award Number UL1TR001412-07. This work is based upon research conducted at the Northeastern Collaborative Access Team beamlines (P30 GM124165, P41 GM103403) utilizing resources of the Advanced Photon Source at the Argonne National Laboratory (DE-AC02-06CH11357). The Eiger 16M detector on 24-ID-E is funded by an NIH-ORIP HEI grant (S10OD021527).

## Author Contributions

T.S.B, J.K.R., and I.K.S. purified and crystallized proteins. T.S.B determined crystal structures.

T.S.B, J.K.R., and I.K.S. performed enzyme inhibition assays. M.J.E supervised the research.

T.S.B, D.E.H, and M.J.E wrote the manuscript. All authors reviewed and approved this manuscript prior to publication.

## Conflicts of Interest

The Eck lab receives (or has received within the past two years) sponsored research support from Novartis, Takeda, and Springworks Therapeutics.

## Methods

### Compounds

All inhibitors were purchased from commercial vendors and are ≥95% pure per vendors’ analyses with the exception of JBJ-04-125-02 which was previously synthesized in-house.^17^ Compound dilutions were prepared in DMSO and stored at −20 °C when not in use.

### Protein expression and purification

The kinase domains of EGFR and HER2, were expressed and purified as previously described.^3,30,31^ Mutations were introduced via PCR and verified by Sanger sequencing. Briefly, insect cells infected with recombinant baculovirus were pelleted and resuspended in buffer, sonicated, clarified via ultracentrifugation, and purified using Ni-NTA agarose beads. Following TEV cleavage of the His-GST tag, the protein was further purified through size exclusion chromatography, concentrated to 3-4 mg/mL, and flash frozen in liquid nitrogen.

### Crystallization and structure determination

Purified EGFR(T790M/V948R) kinase domain was crystallized at approximately 3 mg/mL with 10 mM MgCl_2_ and 1 mM adenylyl-imidodiphosphate (AMP-PNP) via hanging drop vapor diffusion at room temperature. Drops of 1 μL protein were combined with 1 μL well solution consisting of 0.1 M Bis-Tris pH 5.0-6.0 and 20-30% (w/v) PEG 3,350. To obtain the BI-4020 co-crystal structure, single crystals were moved to new 2 μL drops of 0.1 M Bis-Tris and 35% PEG 3,350 over wells of the same composition. Prior to moving crystals, the new drops were supplemented with 0.5 mM BI-4020 (from 10 mM DMSO stock). Crystals were soaked overnight, briefly cryoprotected in well solution containing 20% ethylene glycol, and flash frozen in liquid nitrogen. WT EGFR kinase domain crystals were prepared and soaked as described for T790M/V948R crystals, except the well solution consisted of 0.1 M MES pH 6.5 and 0.8 M sodium citrate and were cryoprotected with 30% ethylene glycol.

Diffraction data were collected at the Advanced Photon Source at the Argonne National Laboratory on NE-CAT beamlines 24-ID-C and 24-ID-E at 100 K. Data were indexed, integrated, and scaled using Dials and Aimless via xia2 compiled by SBGrid.^32–34^ Structures were phased via molecular replacement with PDB 5D41 or 2GS2.^19,35^ Refinement was performed using Phenix with iterative rounds of manual model building in Coot.^36,37^ Custom ligand restraints were generated using eLBOW in Phenix.^38^ Structures have been deposited in the Protein Data Bank (PDB) with the accession codes 7KXZ and 7KY0.

### Kinase inhibition assays

Inhibition assays were performed using the HTRF KinEASE tyrosine kinase assay kit (Cisbio) according to the manufacturer’s protocol. Inhibitors (10 mM DMSO stocks) were dispensed into black 384-well plates using an HP D300e dispenser and normalized to a 1% final DMSO concentration. Assay buffer containing purified EGFR at a final concentration of 5 nM for WT EGFR and HER2, 0.1 nM for L858R, 1 nM for L858R/C797S, and 0.02 nM for L858R/T790M and L858R/T790M/C797S were dispensed using a Multidrop Combi dispenser into 384-well plates and incubated with the inhibitors at room temperature for 30 min. Reactions were initiated with 100 μM ATP using a Multidrop Combi dispenser and allowed to proceed for 30 min at room temperature before being quenched using the detection reagent from the KinEASE assay kit. The FRET signal ratio was measured at 665 and 620 nm using a PHERAstar microplate reader. Data were processed using GraphPad Prism and fit to a three-parameter dose-response model with a Hill slope constrained to −1. The assay was performed three independent times in triplicate.

## Notes

https://www.rcsb.org/structure/7KY0

https://www.rcsb.org/structure/7KXZ

