## Supplemental Table and Figures for "Structural analysis of the macrocyclic inhibitor BI-4020 binding to EGFR kinase"

**SI Table 1:** Data collection and refinement statistics.

| | BI-4020 WT | BI-4020 TM/VR |
|---|---|---|
| PDB Accession | 7KXZ | 7KY0 |
| Resolution range | 51.37 - 2.4 (2.486 - 2.4) | 68.92 - 3.105 (3.216 - 3.105) |
| Space group | I 2 3 | P 1 $2_1$ 1 |
| Unit cell | 145.29 145.29 145.29 90 90 90 | 70.22 100.73 86.79 90 101.07 90 |
| Total reflections | 103473 (10625) | 83605 (7291) |
| Unique reflections | 19877 (1994) | 21014 (2005) |
| Multiplicity | 5.2 (5.3) | 4.0 (3.6) |
| Completeness (%) | 98.86 (99.85) | 97.08 (91.79) |
| Mean I/sigma(I) | 17.57 (2.40) | 3.01 (1.09) |
| Wilson B-factor | 55.57 | 49.08 |
| $R_{merge}$ | 0.0588 (0.4955) | 0.3153 (0.7061) |
| $R_{pim}$ | 0.0278 (0.2342) | 0.1904 (0.4163) |
| $CC_{1/2}$ | 0.998 (0.833) | 0.341 (0.573) |
| Reflections used in refinement | 19876 (1994) | 20950 (1980) |
| Reflections used for R-free | 996 (96) | 1049 (111) |
| $R_{work}$ | 0.197 (0.271) | 0.238 (0.310) |
| $R_{free}$ | 0.214 (0.333) | 0.286 (0.376) |
| Number of non-hydrogen atoms | 2495 | 9502 |
| macromolecules | 2412 | 9342 |
| ligands | 40 | 160 |
| solvent | 43 | 1160 |
| Protein residues | 302 | 0.010 |
| RMS (bonds) | 0.003 | 0.011 |
| RMS (angles) | 0.62 | 1.48 |
| Ramachandran favored (%) | 96.9 | 96.6 |
| Ramachandran allowed (%) | 2.7 | 2.8 |
| Ramachandran outliers (%) | 0.4 | 0.6 |
| Rotamer outliers (%) | 7.6 | 4.0 |
| Average B-factor | 68.3 | 46.4 |
| macromolecules | 68.3 | 46.5 |
| ligands | 75.4 | 37.8 |
| solvent | 60.5 | |

*Statistics for the highest-resolution shell are shown in parentheses.

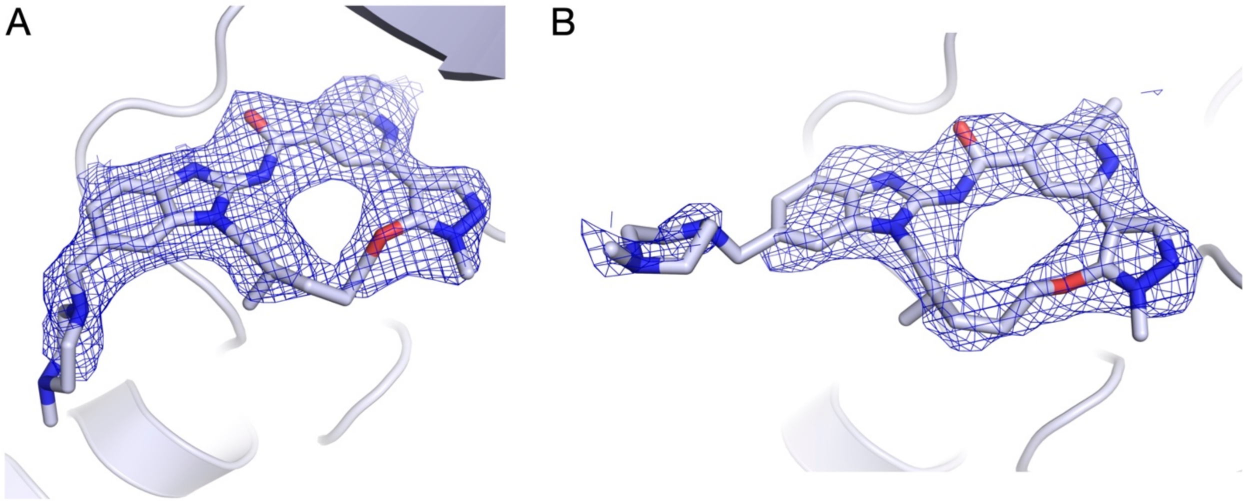


**SI Figure 1: BI-4020** ligand densities. $2F_o$-$F_c$ density contoured at 1 sigma for A) EGFR(T790M/V948R) and B) WT EGFR structures.

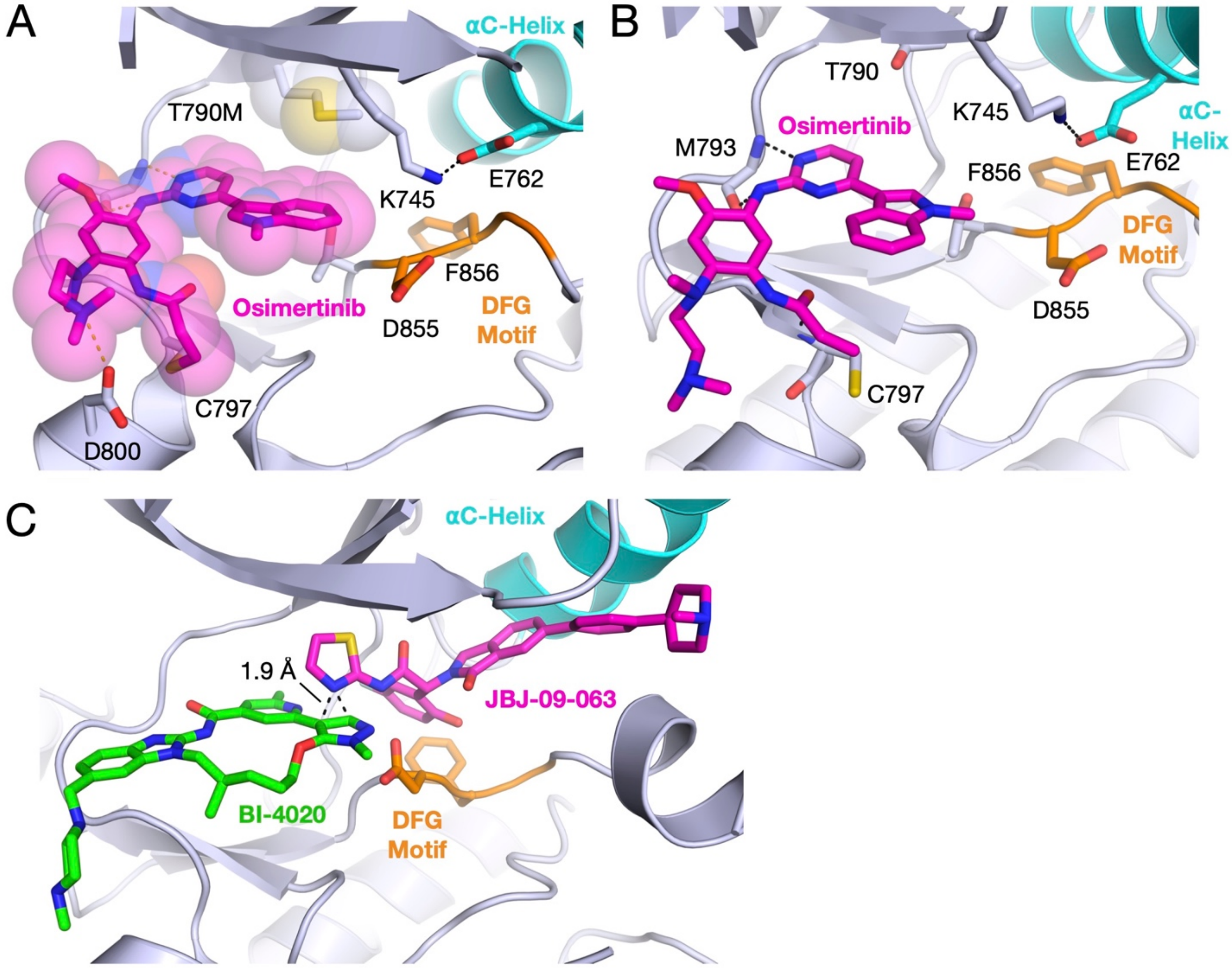


**SI Figure 2:** Ligand comparisons to illustrate binding mode similarities. A) Osimertinib in complex with EGFR(T790M) displaying an unusual flipped indole (compare to panel B) that makes hydrophobic contacts with T790M (PDB 6JX0). EGFR B) Canonical binding mode of osimertinib in complex with WT EGFR (PDB 4ZAU). C) Overlay of BI-4020 and JBJ-09-063 crystal structures (PDB 7KXZ and 7JXQ, respectively). A steric clash precluding co-binding is observed between the inhibitors (distance <2 Å).

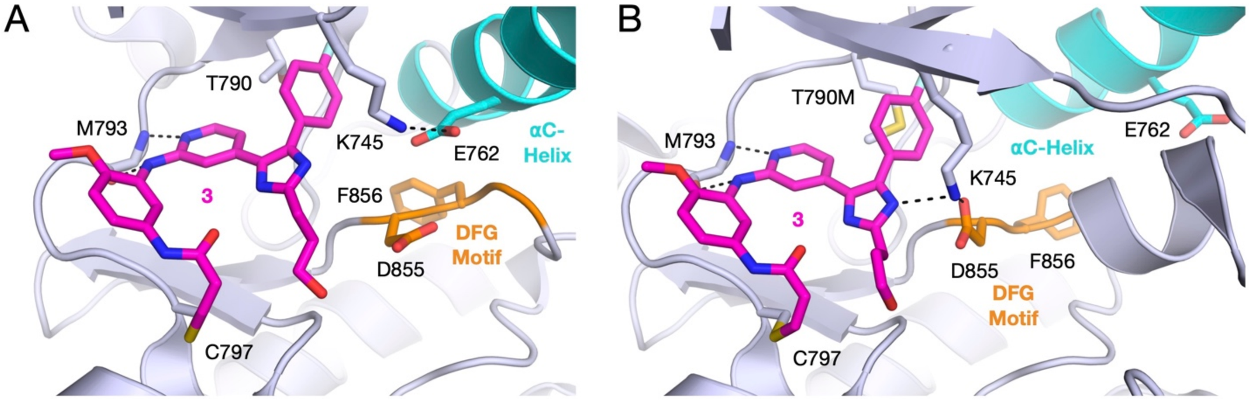


**SI Figure 3:** Conformation of the EGFR kinase domain affects interactions between the trisubstituted imidazole inhibitor **3** (LN2380) and the protein. A) In the active conformation, K745 interacts with E762 (PDB 6VH4). B) In the inactive conformation, the imidazole on **3** interacts with K745 (PDB 6V6O).
